# Thermal selection drives biodiversity origination across the Atlantic/Indian Ocean boundary

**DOI:** 10.1101/419036

**Authors:** Peter R. Teske, Jonathan Sandoval-Castillo, Tirupathi Rao Golla, Arsalan Emami-Khoyi, Mbaye Tine, Sophie von der Heyden, Luciano B. Beheregaray

## Abstract

Intraspecific genetic structure in widely distributed marine species often mirrors the boundaries between temperature-defined bioregions. This suggests that the same thermal gradients that maintain distinct species assemblages also drive the evolution of new biodiversity. Ecological speciation scenarios are often invoked to explain such patterns, but the fact that adaptation is usually only identified when phylogenetic splits are already evident makes it impossible to rule out the alternative scenario of allopatric speciation with subsequent adaptation. We integrated large-scale genomic and environmental datasets along one of the world’s best defined marine thermal gradients (the South African coastline) to test the hypothesis that incipient speciation in the sea is due to divergence linked to the thermal environment. We identified temperature-associated gene regions in a coastal fish species that is spatially homogeneous throughout several temperature-defined biogeographical regions on the basis of selectively neutral markers. Based on these gene regions, the species is divided into geographically distinct regional populations. Importantly, the ranges of these populations are delimited by the same ecological boundaries that define distinct infraspecific genetic lineages in co-distributed marine the species, and biogeographical disjunctions in species assemblages. Our results indicate that ecologically-mediated selection represents an early stage of marine speciation in coastal regions that lack physical dispersal barriers.

## Introduction

Molecular phylogenies of marine species present along continuous coastlines have revealed that spatial disjunctions between distinct evolutionary lineages are often associated with the boundaries between different marine biogeographic regions [1,2], but such genetic patterns tend to be present in only a fraction of species [1,3,–6] (Fig. 1). This discrepancy is often attributed to life history: actively dispersing species, and those with extended planktonic dispersal phases, cross the boundaries between bioregions more frequently than species with short propagule duration, making them less likely to diverge in spatial isolation [5,7]. However, support for this paradigm is not consistent, as numerous studies from North America [6,8], South Africa [1,9] and Australasia [4,10] have failed to identify a clear link between genetic structure and dispersal potential.

**Figure 1.**
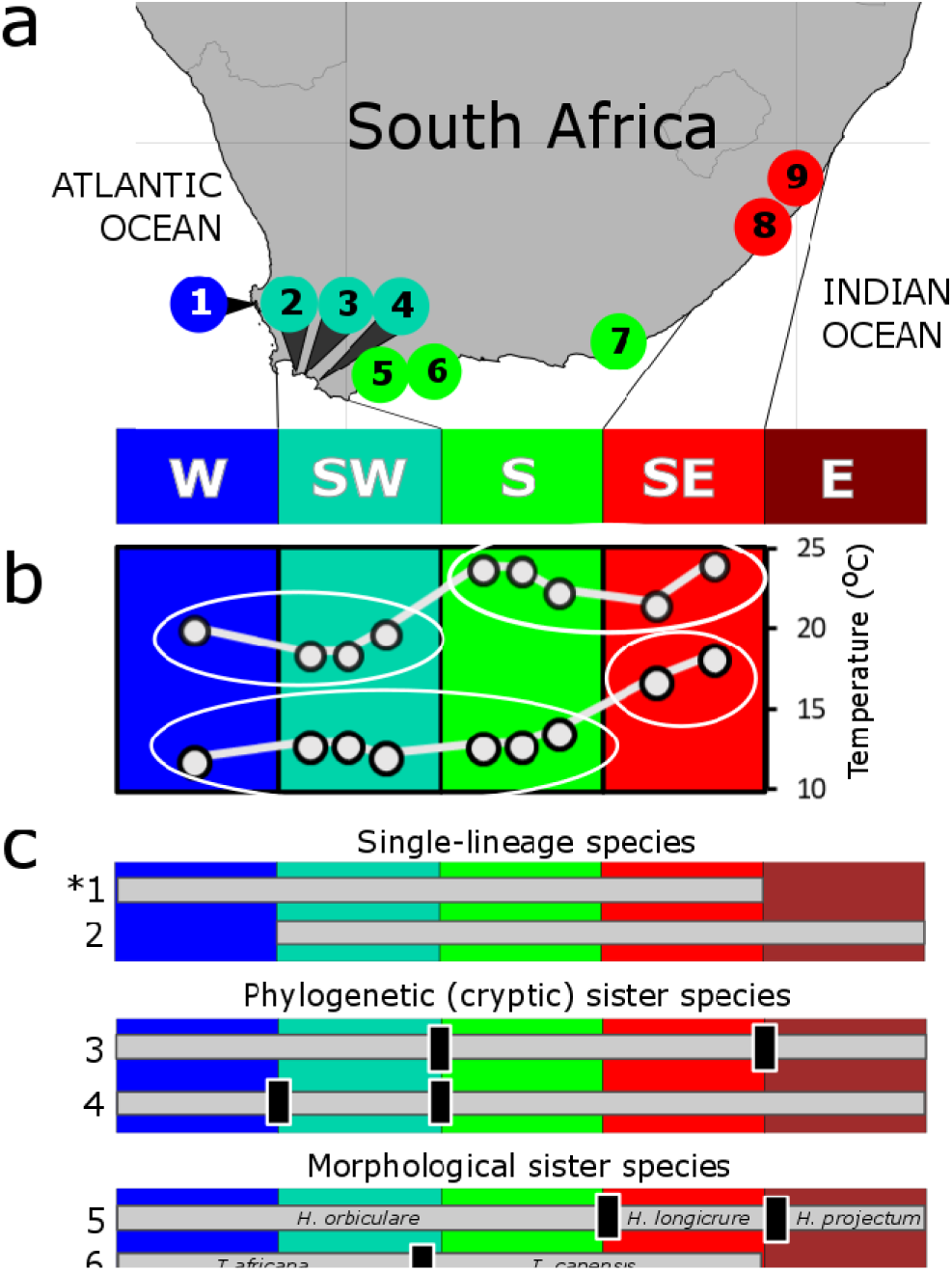
Sampling sites, marine bioregions and examples of genetic breaks in South African coastal animals; **a** A map indicating the location of sampling sites within southern Africa’s temperature-defined marine bioregions; **b** maximum and minimum sea surface temperatures at each sampling site; these temperatures divide bioregions into two groups each (indicated by ellipses), and by themselves only partially explain the region’s biogeography; **c** examples of distribution ranges (grey horizontal bars) and location of genetic breaks (black vertical bars) in coastal South African animals, arranged hierarchically. The top panel depicts species that occur as a single phylogenetic lineage in multiple bioregions: 1. *Psammogobius knysnaensis* (the study species, marked with an asterisk) [24] and 2. *Scutellastra longicosta* [16]. The middle panel depicts species with phylogenetically distinct sister lineages that are not distinguishable morphologically (cryptic species): 3. *Callichirus kraussi* [25] and 4. *Palaemon peringueyi* [26]. The bottom panel depicts morphologically distinguishable sister species: 5. *Hymenosoma* spp. [27] and 6. *Tricolia* spp. [28]. Abbreviations: W, cool-temperate west coast; SW, transition zone on the south-west coast; S, warm-temperate south coast; SE, transition zone on the south-east coast; E, subtropical east coast.

An alternative explanation for this paradox is offered by ecological divergence that preceded the allopatric distribution patterns evident on the basis of selectively neutral genetic markers. This is primarily supported by ‘phylogenetic shifts approaches’, in which phylogenetic splits coincide with ecological divergence [11]. The evidence for ecological speciation is particularly strong when phylogenetic splits are not associated with physical dispersal barriers that can completely isolate sister lineages [11], when contact zones are located in regions where environmental conditions are intermediate [1,12] and when each lineage displays reduced fitness in the habitat of its sister lineage [13–15]. Phylogenetic splits that are shared by multiple species across the same boundary may differ considerably in age [1,16], and by extension, this supports the hypothesis that species in which phylogenetic divergence is not yet evident have undergone ecological differentiation very recently.

The phylogenetic evidence for ecological drivers of speciation is nonetheless circumstantial, because divergence events mostly occurred during the Pleistocene or earlier [16–18], and it is difficult to extrapolate from contemporary conditions when species’ historical distribution patterns are unknown and past oceanographic conditions not well understood. Because of such uncertainties, it is controversial to ascertain whether adaptation to divergent environments that reduced levels of gene flow because of the maladaptation of migrants was the primary driver of divergence, or whether it occurred after a phylogenetic split that may very well have evolved during an extended period of physical isolation.

More compelling evidence for ecological speciation in the sea would come from scenarios in which there is support for genetic differentiation that coincides with biogeography, but in which phylogenetic divergence indicative of speciation has not yet occurred [15]. The fact that phylogeographic breaks tend to be present in only a fraction of the species whose ranges span the boundaries between ecologically distinct marine regions [1,3,4] suggests that the condition of recent divergence may be met by those species that display no genetic divergence on the basis of the selectively neutral datasets typically employed in phylogeographic studies [19,20].

The South African coastline is characterised by ecologically distinct marine bioregions (Fig. 1) that are arranged along a thermal gradient [1]. This provides a unique opportunity for studying the importance of incipient environmentally-driven parapatric speciation in the sea, as biogeography (and, by extension, ecological speciation) is believed to be primarily a function of species’ thermal tolerance ranges [21–23]. Numerous species complexes exist along this coastline that comprise cryptic species whose ranges are limited by the boundaries between bioregions [1], and which exhibit distinct temperature preferences [13,14]. This suggests that thermal adaptation contributes towards limiting gene flow between biogeographic regions by reducing migrant fitness and by subjecting migrants to competitive exclusion [1]. However, in some species, a single evolutionary lineage is found across multiple bioregions (Fig. 1). The latter are suitable candidates for determining whether diversifying selection driven by the environment, and corresponding reductions in gene flow, may have preceded phylogenetic splits.

We tested this hypothesis by generating genome-wide data from one of these phylogenetically homogeneous species [24], the Knysna sandgoby, *Psammogobius knysnaensis* (Fig. 1). We expect population divergence that mirrors coastal biogeography to be evident based only on temperature-associated genes. This would present compelling evidence that in coastal regions that lack physical dispersal barriers, thermal selection plays a defining role in the early stages of parapatric ecological speciation.

## Methods

### (a) Sampling procedure

Tissue samples from a total of 312 individuals of *Psammogobius knysnaensis* were collected from the mouth areas of nine estuaries throughout the species’ range (Table 1) using a pushnet. Upon capture, a fin clip was obtained from one of the pectoral fins using sterilised fingernail scissors, and immediately preserved in 100% ethanol. The fish subsequently released. Samples in the West were Coast National Park (Langebaan Lagoon) were collected under research permit no. CRC/2015/033--2015/V1.

### (b) Generation and processing of genomic data

Genomic DNA was extracted using the CTAB protocol [29], and double digest restriction site–associated DNA (ddRAD) libraries were constructed for a subset of 129 individuals and 12 replicates with particularly high quality DNA, following the protocol described in [30] and modified as described in Sandoval-Castillo et al. [31]. Libraries were pooled in groups of 48 or 93 samples per lane and sequenced on an Illumina HiSeq 2000 (100 bp paired-end reads) platform at the McGill University and Genome Québec Innovation Centre. Raw sequences were processed as described in the Supporting Information.

### (c) Identification of loci under thermal selection and neutral loci

We assessed the contribution of coastal sea surface temperature (SST) to the overall pattern of genetic differentiation using the R package gINLAnd [32]. This software uses a spatial generalized linear mixed model to quantify the correlation between genotypes and environmental variables, while controlling for the effects of spatial population structure and population history. Briefly, gINLAnd estimates the covariance associated with the spatial distribution of the samples and a locus-specific effect of each environment variable; it then estimates the likelihood of two competing models: a model with the environmental effect and a reduced model without the environmental effect. Finally, gINLAnd assesses the strength of genetic dependence on the environmental variab e by computing a Bayes factor between the two models. To avoid false positives, we used a conservative approach in which only those loci which showed a log Bayes factor (BF) ≥10 were considered to be under selection (a log BF>4.6 is considered decisive [33]). A plot depicting loci under selection is shown in Fig. S1. We calculated a multidimensional scaling projection of the coastal distance between sampling sites using the R package MASS 7.3 [34], which is more meaningful than using the original geographic coordinates because this would have required connecting sites via terrestrial habitat. As the application of satellite-based SST data is often problematic when studying coastal biogeography, because it includes data from offshore regions [35], we used southern African temperature data based on *in situ* measurements, as described in the Supporting Information.

To compare the temperature-associated loci with a data set comprising only selectively neutral loci, the following approach was used. Thermal selection may be only one of a number of drivers of selection, so we used BayeScan v. 2.1 [36] to identify markers under selection on the basis of outlier scans rather than temperature data. This method was used because it has a low error rate compared to other tests for the detection of outlier loci [37]. Default settings were applied with prior odds set to 10, but a very high false discovery rate of 20% was applied to create a neutral data set with a low probability of containing any remaining loci under selection. We then excluded 304 outlier loci, together with 27 additional loci identified by gINLAnd that were not found by BayeScan, to create a data set of 8201 selectively neutral loci.

### (d) Functional annotation

To identify the possible functions of genes under thermal selection, we blasted the flanking sequences of temperature-associated loci against the NCBI non-redundant nucleotide database. The resulting reads were then annotated against the UniProtKB/Swiss-Prot database [38]. We then performed a gene ontology term analysis in topGO 2.24.0 [39]. Genes whose function indicates an influence of thermal selection were identified by searching the relevant literature.

### (e) Population genetic structure

Genetic structure was investigated separately for loci under selection and neutral loci. We employed both clustering and phylogenetic approaches. Discriminant Analysis of Principal Components (DAPC) was performed with the R package ADEGENET v. 2.1.0 [40]. DAPC defines a model with synthetic variables in which the genetic variation is maximized between clusters of individuals (*K*), and minimized within clusters. We used *k*-means clustering and the Bayesian Information Criterion (BIC) to identify the best-supported number of clusters. Patterns of genetic structure were also explored using *fastStructure* 1.0 [41], which uses variational Bayesian inference under a model assuming Hardy-Weinberg equilibrium and linkage equilibrium. We used a simple prior and set all other parameters to the default value, expect for the convergence criterion, which was lowered to 10^-8^. The programme was run for each value of *K* = 1-9 independently, and each value was cross-validated 1000 times. The python script *chooseK* was used to identify an optimal range of *K* values, and the resulting barplots were visualised with the R package distruct2.2 [42].

Phylogenetic analyses were performed in BEAST v. 2.4.7 [43]. A maximum clade credibility (MCC) tree was reconstructed using a discrete phylogeographic analysis [44]. In this case, the data set comprised individual alleles of each individual that were reconstructed in PHASE v. 2.1.1 [45] using default settings. When more than one pair of haplotypes was possible for an individual, the one with the highest probability was used. In addition to reconstructing a phylogenetic tree, this method can infer the most likely bioregion in which each ancestral node in the MCC tree was present. One hundred million generations were specified, and trees saved every 100 000 generations, and the first 20% of trees were discarded as burn-in. Model and prior settings followed those recommended in the tutorial available at http://hpc.ilri.cgiar.org. For comparison, a corresponding MCC tree was created for previously published mtDNA COI data [24] using the same settings.

## Results

The ddRADseq [30] procedure was used to generate a genome-wide dataset of single nucleotide polymorphisms (SNPs) from individuals collected at nine sites that are located within four temperature-defined marine bioregions (Fig. 1). A total of 405,648,596 raw reads were generated on two Illumina lanes. After demultiplexing and quality filtering, an average of 1,560,510 reads were obtained per individual, totalling 224,713,440 reads. The filtered catalogue resulted in 8,532 ddRADseq loci containing 15,633 SNPs. A final dataset was obtained by extracting only the SNPs with the best quality score from each polymorphic ddRADseq locus to remove SNPs that are likely in linkage disequilibrium. After removing individuals with more than 20% missing data, the final data set comprised 109 individuals genotyped for 8,532 SNPs. We then used a spatially explicit generalized linear mixed model to test for direct associations between SNP allele frequencies and temperature-related variables, while controlling for the effects of spatial structure and shared population history, using the program gINLAnd [32]. Unlike *F*_ST_- based outlier scans, which identify loci on the basis of population information [36], the identification of loci in in genotype-environment association methods such as gINLAnd is thus not influenced by any regional population structure [46]. We explored various combinations of maximum or minimum temperature as the environmental variable with covariance factors that included geographic distance, biogeographic boundaries and a combination of the two. A simplistic resistance matrix approach [47] was used, where geographic distance and biogeographic boundaries between pairs of sites were given a resistance value (one unit of resistance per km between sites and ten units of resistance if sites were located in different marine bioregions). We then calculated multidimensional scaling projections based on the resistance pairwise matrices, and these were used as covariance factor to control for spatial structure. We further explored the effect of using only SNP data from the coolest and the warmest marine bioregions, using geographic distance as the controlling factor (see Table S1 for detail on number of SNPs identified). Most subsequent analyses were performed with the data set recovered using minimum temperature (with geographic distance as the covariance factor), which is thought to represent a key selective agent of adaptive divergence in the study region because it can limit species’ distributions [48,49].

SNPs from ddRADseq originate from all genomic regions, and some may be located on protein-coding genes that are strongly affected by temperature. While such associations may not necessarily imply a causal relationship, identifying their function may contribute towards an improved understanding of possible drivers of genetic divergence between temperature-defined bioregions. Although no fully annotated transcriptome for the family Gobiidae is presently available, nine of the loci (identified using either maximum or minimum temperature, with geographic distance as the covariance factor) could be annotated as genes involved in mitigating thermal stress (Table S2). Three of these (14-3-3 gene, tyrosine protein kinase and tubulin beta chain) are of particularly interest because they were involved in heat stress responses in a species of goby [50], or cold stress adaptation/acclimation in other teleosts [51,52]. In all but two cases, loci that were identified using minimum temperature data were also identified using maximum temperatures (Table S2). This suggests that even though most experimental studies investigated responses to heat stress, genetically fixed differences of these genes between temperature-defined marine bioregions may reflect general adaptations to different thermal environments, and thus play a role in determining thermal tolerance ranges.

To ascertain that the study species does not yet exhibit genetic divergence based on putatively neutral data, which could indicate that geographical isolation preceded thermal adaptation, we created a reduced dataset comprising selectively neutral data. In addition to excluding loci identified as being under thermal selection, we excluded outlier loci from genome scans [36] to better estimate demographic parameters and population differentiation [31].

Two complementary methods of investigating a link between population structure and biogeography were performed on both the temperature-associated loci and the neutral loci, Discriminant Analysis of Principal Components (DAPC) [53] and fastStructure [41]. Statistical support was high for 3-4 clusters (*K*) when analysing the temperature-associated loci, while a single cluster (*K* = 1) was found for the neutral loci with both methods (Fig. S2, Fig. S3). Even though temperature alone only accounts for two marine bioregions (Fig. 1), and despite the fact that geographic distances and/or the boundaries between bioregions were controlled for when identifying temperature-associated loci, an affiliation of genetic clusters with up to four bioregions was found when using the temperature-associated loci. The distinctness of subtropical (SE) individuals from the temperate (W, SW and S) sites was evident in all analyses, and most analyses using minimum temperature as the environmental variable also identified the W coast as a distinct cluster. There was even evidence for distinct SW and S coast clusters, although these were comparatively poorly differentiated. This result was robust, and also recovered using fastStructure (Fig. 2).

**Figure 2.**
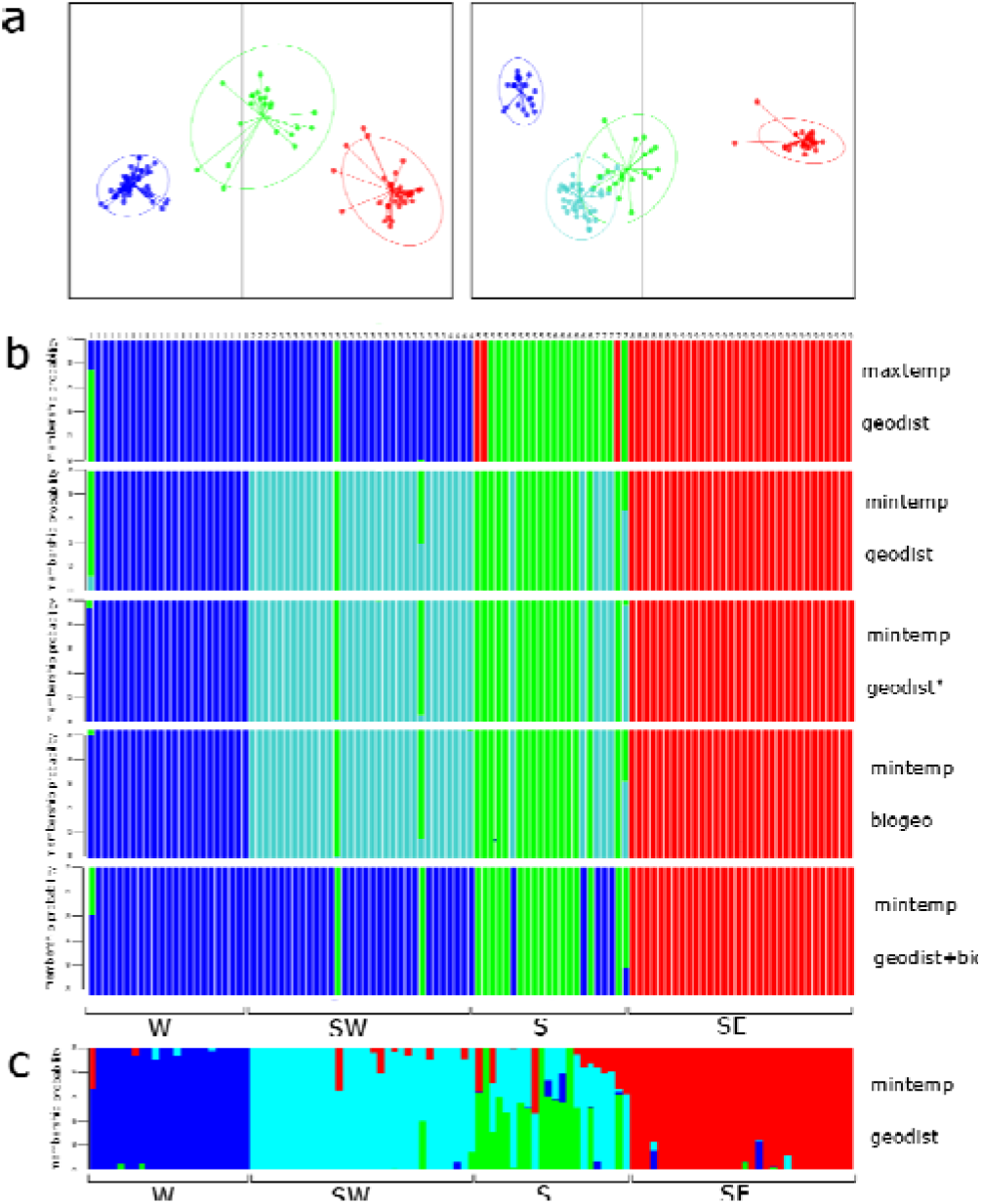
Population genetic structure inferred for temperature-associated loci. **a** DAPC scatterplot with inertia ellipses representing 95% confidence intervals, with colours reflecting the dominant bioregion represented in a particular cluster (left: loci correlated with maximum temperature; right: loci correlated with minimum temperature, in both cases controlling for geographic distance); **b** DAPC compoplots indicating membership probabilities for each individual (vertical bars) within one of four genetic clusters; correlation with temperature and the controlling factor are indicated on the right (maxtemp = maximum temperature, mintemp = minimum temperature, geodist = geographic distance, biogeo = biogeography; *indicates that only sites 1, 8 and 9 were used to find loci correlated with minimum temperature); **c** corresponding concensus fastStructure barplot for four genetic clusters (*K*); for comparison, barplots for *K*=2-5 are shown in Fig. S4. Site numbers and abbreviations correspond to those in Fig. 1 and Table S3.

Congruent with the clustering methods, a maximum-clade credibility tree (Fig. 3a) of temperature-associated loci (minimum temperature with geographic distance as the covariance factor) recovered both the western and south-eastern group as mostly distinct but poorly differentiated clusters nested within a tree whose oldest nodes were inferred to have existed on the south coast. Some branches are nested within clades that mostly have location states from other regions, which may reflect migration between adjacent marine bioregions. For comparison, a maximum-clade credibility tree reconstructed from mtDNA COI sequences [24] shows no clear regional structure (Fig. 3b).

**Figure 3.**
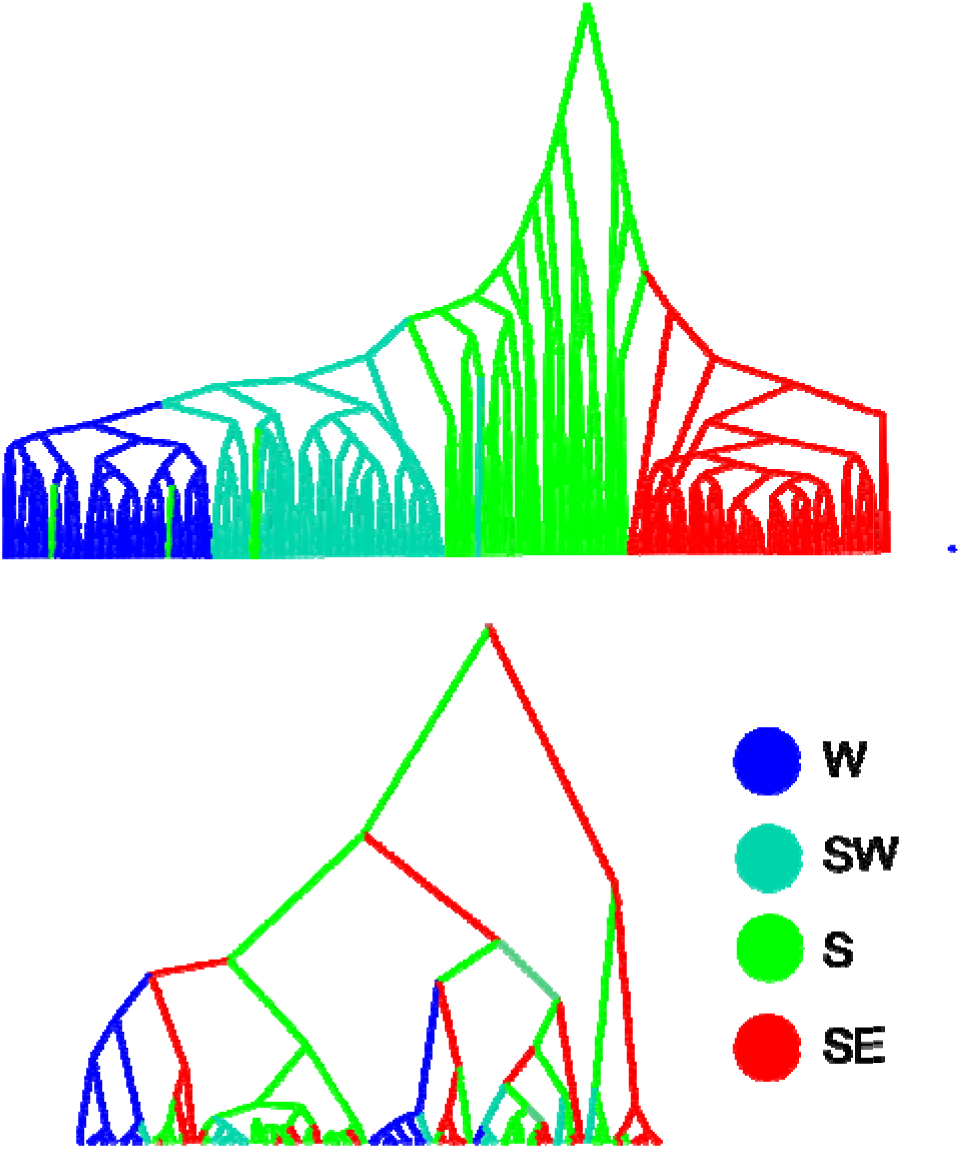
Reconstruction of phylogenetic relationships between individuals of *Psammogobius knysnaensis* from four South African marine bioregions using maximum-clade credibility trees from **a** phased SNP data of temperature-associated loci (correlated with minimum temperature) and **b** mtDNA COI data, with location state reconstructions of ancestral nodes. In **a**, clear regional structure is evident, but there are possible migrants (some branches are nested within clades that mostly have location states from other regions). In contrast, there are no clear regional clades in **b**. Site numbers and abbreviations correspond to those in Fig. 1 and Table 1, and trees are not drawn to scale.

## Discussion

Speciation is a continuous process comprising a number of evolutionary stages that range from adaptive differentiation to complete reproductive isolation between populations [54]. Identifying the primary drivers of speciation is challenging because a considerable amount of time has often already passed by the time incipient speciation becomes evident. This makes it difficult to distinguish ecologically-driven divergence with ongoing gene flow from allopatric divergence and secondary contact [55,56].

Biogeography is often considered to be a function of species’ thermal tolerance ranges [21–23]. The fact that South Africa’s coastal biogeography is mirrored by intraspecific spatial genetic structure suggests that species present in more than one province should comprise multiple evolutionary lineages that represent cryptic species [1]. The goby *Psammogobius knysnaensis* is one of a number of coastal southern African species that occur in multiple marine bioregions, but which displays no regional divergence on the basis of selectively neutral markers [24] of the type that are primarily used in phylogeographical research [19,20]. Here, we reject the previous finding of genetic homogeneity and show that this species is in fact represented by multiple regional groups delimited by temperature-defined bioregions. The fact that this is only evident for temperature-associated loci, and not for putatively neutral loci, confirms that divergence must have taken place in the absence of an interruption of gene flow due to physical dispersal barriers. Under these conditions, a pattern of isolation-by-adaptation [57] can be expected to eventually evolve, as migrants dispersing into adjacent bioregions will have fewer surviving offspring and reduced survival rates compared to residents. A west-to-east thermal differentiation was evident particularly for the minimum temperatures, where the two easternmost sites had much warmer water than the other sites (Fig. 1). However, there was no indication that marine bioregions could be identified on the basis of high or low temperatures alone, and the identification of loci under thermal selection and subsequent detection of up to four genetic clusters cannot be explained as being an artefact of the temperature variables used in the gINLAnd analyses.

Adaptations to the thermal environment are complex and ubiquitous in nature. Temperature affects many different biological pathways, with strong effects on the integrity of proteins and cellular structures and on the rates of physiological processes, particularly in ectotherms [58,59]. The thermal environment can promote partial reproductive isolation between populations, which might drive them along the speciation continuum [60]. This is particularly true for organisms that (i) have distinct populations with parapatric distributions along the thermal gradient, (ii) do not maintain a stable internal temperature (poikilotherms), and (iii) are found across stable thermal gradients (e.g. aquatic environments), which are regions where exogenous divergent selection is not expected to weaken due to marked temperature fluctuations [60]. Our study system meets all these conditions and represents an example of parapatric ecological divergence with genomic hallmarks of incipient evolutionary divergence driven by the thermal environment.

Unlike previous spatial demographic inferences from coastal southern Africa, which typically reflect the influence of past climatic changes [16,17,61,62], the spatial genetic patterns identified here can be explained by present-day environmental conditions. On the east coast, northward dispersal in the nearshore area is facilitated by wind-driven circulation [63], but this is unlikely to occur beyond site 9 (the northern distribution limit of *P. knysnaensis*) [62,64] because under contemporary conditions, the southward-flowing Agulhas Current flows very close to the coast and causes the parallel southward flow of nearshore circulation [65]. In the western portion of the species’ range, gene flow between south and west coast is primarily facilitated by the westward drift of surface water [66]. The limited evidence for gene flow in both cases would be difficult to explain if one exclusively invoked physical isolation, given the high dispersal potential of the species’ larvae coupled with the region’s strong ocean circulation. It suggests that migrants from a particular bioregion are maladapted to the environmental conditions in adjacent bioregions. For example, the distinctness of the west coast population from those on the south-west and south coast may reflect the influence of cold-water upwelling in the west [67].

We hypothesise that thermal selection, perhaps in combination with factors such as oceanography and primary productivity that covary with temperature to influence local adaptation [60], acts primarily on the sensitive larvae. Under this scenario, ecologically diverging populations are limited in their ability to exchange genes and, as such, reproductive isolation is expected to ensue [60]. There are no known conspicuous phenotypes that differ between the presumably locally-adapted *P. knysnaensis* populations, but this is unsurprising because thermal adaptation often initially creates cryptic changes at the level of cell membranes or thermal stability of enzymes [68]. Studies that combine information from population genomics and controlled laboratory experiments using temperature-defined populations along an evolutionary continuum of speciation are expected to improve the identification of phenotypes enriched for selection signals of thermal adaptation.

## Conclusion

Allopatric speciation in the marine environment is often invoked along continuous but ecologically subdivided coastlines, despite evidence that the physical dispersal barriers to whom this is attributed are insufficient to completely isolate regional populations [69–71]. Our study contributes to the growing evidence that in adjacent, temperature-defined marine provinces, divergence of loci linked to the thermal environment can precede significant spatial divergence of selectively neutral markers [72,73]. This strongly favours a scenario of parapatric ecological divergence over one in which allopatric divergence is followed by thermal adaptation. In the context of larger biogeographical patterns, where range boundaries in the sea often coincide with the boundaries between temperature-defined bioregions [74,75], this evidence suggests that temperature-driven diversifying selection may be an important early-stage factor in the evolution of marine biodiversity.

## Authors’ contributions

P.R.T. and L.B.B. designed the study, P.R.T., T.R.G. and S.v.d.H. collected the samples, J.S.-C. conducted the experiments, J.S.-C., P.R.T., M.T., A.E.-K. and T.R.G. analysed the data, P.R.T. and L.B.B. wrote the paper, with input from S.v.d.H. and J.S.-C.

## Acknowledgements.

We are grateful to Robert Schlegel and Albertus Smit for providing the SST data. This study was funded by the PADI Foundation (Grant No. 10981 to PRT), the National Research Foundation (CSUR Grant No. 87702 to PRT), the University of Johannesburg (URC/FRC grant to PRT) and the Australian Research Council (FT130101068 and DP110101275 to L.B.B.). The authors are grateful to the Centre for High Performance Computing (CHPC), particularly Dane Kennedy, for supercomputer resources and bioinformatics support. M.T., T.R.G. and A.E.-K. acknowledge the University of Johannesburg for Global Excellence and Stature (GES) fellowships.

